# Blocking the Rac1-TRPC5 pathway protects human podocytes

**DOI:** 10.1101/2020.08.28.272344

**Authors:** Yiming Zhou, Choah Kim, Juan Lorenzo B. Pablo, Fan Zhang, Ji Yong Jung, Li Xiao, Silvana Bazua, Maheswarareddy Emani, Corey Hopkins, Astrid Weins, Anna Greka

**Affiliations:** Department of Medicine, Brigham and Women’s Hospital and Harvard Medical School, Boston, MA, USA; Broad Institute of MIT and Harvard, Cambridge, MA, USA; Department of Internal Medicine, Gachon University Gil Medical Center, College of Medicine, Incheon, Republic of Korea; Department of Pharmaceutical Sciences, University of Nebraska Medical Center, Omaha, NE, USA; Department of Pathology, Brigham and Women’s Hospital, Harvard Medical School, Boston, MA, USA; Medical Research Center, Sun Yat-sen Memorial Hospital, Sun Yat-sen University, Guangzhou, Guangdong, China

## Abstract

Podocyte injury and the appearance of proteinuria are key features of several progressive kidney diseases. Genetic deletion or selective inhibition of TRPC5 channels with small-molecule inhibitors protects podocytes in rodent models of disease, but less is known about the human relevance and translatability of TRPC5 inhibition. Here, we investigate the effect of TRPC5 inhibition in puromycin aminonucleoside (PAN)-treated human iPSC-derived podocytes and kidney organoids. We first established that systemic administration of the TRPC5-specific blocker AC1903 was sufficient to protect podocyte cytoskeletal proteins and suppress proteinuria in PAN-induced nephrosis in rats, an established model of podocyte injury and progressive kidney disease. PAN treatment also triggered the Rac1-TRPC5 injury pathway in human iPSC-derived podocytes and kidney organoids. TRPC5 current was recorded in human iPSC-derived podocytes, and was blocked by AC1903. The TRPC5 blocker also reversed the effects of PAN-induced injury in human podocytes in both 2D and 3D culture systems. Taken together, these results revealed the relevance of the TRPC5-Rac1 pathway in human kidney tissue highlighting the potential of this therapeutic strategy for patients.

## Introduction

Progressive chronic kidney disease (CKD) is associated with increased risk of kidney failure^1^, and its prevalence is rapidly increasing with now more than 850 million people with CKD worldwide^2^. Despite these rising numbers, the therapeutic options available to slow or prevent disease progression are limited^3,4^. Nephrotic syndrome is an important driver of CKD. Characterized by the presence of large amounts of albumin spilling into the urine, nephrotic syndrome is the consequence of damage to the filtering unit of the kidney, the glomerulus. When intact, the kidney filter, made up of endothelial cells, the basement membrane, and the podocytes, is essential for retaining proteins in the blood and removing waste from the body. Many chronic kidney diseases are associated with the loss of podocytes, critical post-mitotic, terminally differentiated cells of the kidney filter that cannot be renewed once lost^5-9^. Due to their limited capacity to proliferate, podocytes are especially vulnerable to various stimuli that lead to injury^10^. Preventing podocyte injury therefore remains a critical objective for the development of effective, targeted therapeutic strategies for kidney diseases.

Numerous studies indicate that dysfunction of the podocyte cytoskeleton contributes to progressive proteinuric kidney diseases ^4, 11^ such as Focal Segmental Glomerulosclerosis (FSGS) and Minimal Change Disease (MCD). Decreased expression of podocyte cytoskeletal proteins, including synaptopodin, nephrin, and podocin, is an early event in podocyte injury that results in disorganization of the cytoskeleton, the fusion of foot processes, and ultimately the development of proteinuria and subsequent kidney damage^12^. A significant number of mutations associated with filter barrier damage result in excess Rac1 signaling in podocytes including mutations in *ARHGAP24*^*13*^, *ARHGDIA*^*14*^, and *ARHGEF17*^*15*^. A small GTP-binding protein, Rac1 is closely associated with various proteinuric kidney diseases, and critically, the regulation of podocyte cytoskeletal proteins. In addition to disruption of cytoskeletal protein remodeling^16^, Rac1 activation results in increased ROS production and regulation of ion channels.

Ion channels are critical to kidney function, and their involvement in kidney disease is an active area of investigation. Transient receptor potential (TRP) channels are receptor-operated, nonselective, Ca^2+^-permeable, cationic channels that were first identified in *Drosophila*^*17, 18*^. TRPC (TRP canonical) channels are a subgroup of this larger family that are particularly relevant to podocyte biology^19^ and have been shown to play an important role in the pathogenesis of kidney disease. Ca^2+^ influx^20^ through TRPC5 elicits dynamic and tightly regulated biochemical responses that activate Rac1. Rac1 activation leads to further vesicular insertion of TRPC5 into the plasma membrane thus making more TRPC5 channels available for activation and completing a feed-forward pathway. Critically, data from three chemically distinct compounds that block TRPC5 activity (AC1903, ML204, and GFB-8438) have demonstrated beneficial effects when applied to rodent models of kidney disease^21^. In addition to TRPC5, both gain-of-function and loss-of-function mutations in TRPC6 channel activity contribute to podocyte injury^22^ further implicating TRPC channel activity in chronic kidney diseases.

While the role of TRPC5 in a Rac1-TRPC5 feedforward injury circuit in podocytes has been defined, whether TRPC5 activity drives disease-relevant phenotypes in human kidney cells remains unexplored. The current study addresses these questions directly by harnessing the technological advances afforded by human induced pluripotent stem cell (iPSC)-derived 2D podocyte cultures (iPodos) and 3D kidney organoids. We determined that human podocytes express functional TRPC5 channels, and that TRPC5 inhibition protects human podocytes from injury. Our data were cross-validated in the experimentally tractable PAN-induced nephrotic rat model. This work provides a strong rationale for ongoing efforts to move TRPC5 inhibitors into the clinic (NCT03970122; https://clinicaltrials.gov/) for the treatment of progressive proteinuric kidney diseases.

## Results

### Inhibition of TRPC5 channel activity reduces proteinuria and protects podocytes from injury in PAN-treated rats

PAN injection administered to rats causes progressive renal injury that resembles MCD or FSGS. The extent of damage depends on the amount and frequency of the PAN injection^23, 24^. Many molecules, including TRPC6 channels, are associated with PAN-induced nephrosis in rats^25-28^. However, a recent study showed little to no protective effects in the early phase of PAN treatment in rats with genetic deletion of TRPC6 channels^29^, suggesting that other pathways may mediate early stage disease. Previously, we have shown that inhibition of TRPC5 protects podocytes from injury and loss in the early phases of disease in several rodent models, suggesting a clinically relevant role for TRPC5 inhibition.

To investigate the role of TRPC5 in PAN nephrosis, we administered a single dose of PAN (50 mg/kg body weight) to rats, which induced a significant amount of urine albumin 7 days after injection. In contrast, co-administration of the TRPC5 channel inhibitor AC1903 twice per day significantly reduced urine albumin 7 days after PAN injection (Fig. 1A). Hemotoxylin and eosin staining showed no obvious morphological changes in glomeruli and tubules from all groups (Fig. 1B); however, transmission electron microscopy showed extensive foot process effacement (FPE) without changes to the glomerular basement membrane (GBM) or the mesangial cells (Fig. 1C), resembling the clinical manifestations of MCD in patients. Statistical analysis of rat podocyte foot processes (FPs) showed that treatment with t AC1903 preserved FP number and protected FPs from effacement (Fig. 1D and E). These results indicate that TRPC5 channels *in vivo* play an important role in inducing injury by PAN.

**Figure 1.**
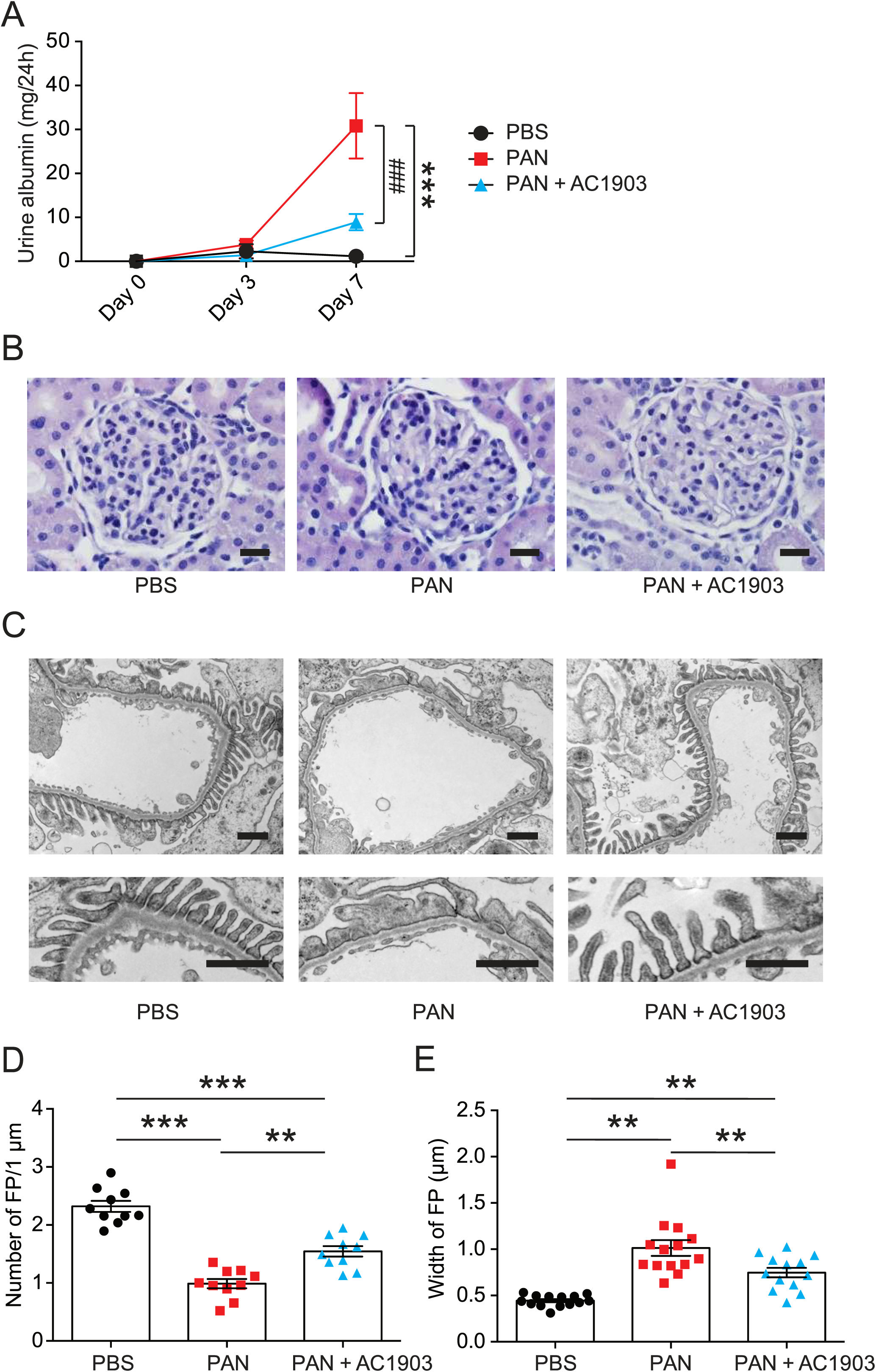
AC1903 reduces proteinuria and protects podocytes from injury in a PAN nephrosis rat model. (**A**) 24-hour urine albumin levels from PBS, PAN and PAN + AC1903 treated rats on day 0, 3 and 7. PAN 50 mg/kg, AC1903 50 mg/kg. PBS n = 6; PAN n = 15; PAN + AC1903 n = 13. ***p < 0.001, ###p < 0.001. (**B**) Representative PAS staining images of PBS, PAN and PAN + AC1903 treated rats on day 7. Scale bar 20 µm. (**C**) Representative TEM images of podocyte foot processes (FPs) from PBS, PAN and PAN + AC1903 treated rat on day 7. Scale bar 1 µm. (**D, E**) Quantification of podocyte FPEs using the FP number (**D**) and width (**E**) on 1 µm glomerular basement membrane from PBS, PAN and PAN + AC1903 treated rats on day 7. *p < 0.05, ***p < 0.001.

We further characterized the effects of PAN injection on several important podocyte proteins. In PAN-induced nephrosis rats, the abundance of two podocyte cytoskeletal proteins, podocin and synaptopodin, was reduced in PAN-treated rat kidneys, while the expression levels of podocyte transcription factor WT1 were not affected, indicating that PAN at this concentration caused alterations in podocyte cytoskeletal structure but did not drive cell loss. Treatment with the AC1903 successfully restored the PAN-induced depletion of podocin and synaptopodin (Supplementary Figure 1). Thus, we concluded that inhibition of TRPC5 channel activity can reduce FPE by protecting podocyte cytoskeletal structure.

### Systemic administration of TRPC5 inhibitor reduces PAN-induced glomerular channel activity

To understand TRPC5 channel involvement and contribution to PAN-induced podocyte injury, we performed TRPC5 single-channel recordings from acutely isolated rat kidney glomeruli according to our previously reported protocol and procedures^16^. A single dose of PAN treatment successfully increased TRPC5 single channel activity in response to the TRPC5 agonist riluzole, while systemic co-administration of AC1903 with PAN in rats significantly reduced TRPC5 activity from isolated glomeruli (Fig. 2A). PAN-treated rats showed a higher NPo value, the product of channel number and open-channel probability, while AC1903-treated rats exhibited a lower NPo value (Fig. 2B). We hypothesized that systemic AC1903 administration would significantly lower the number of TRPC5 channels inserted in the podocyte plasma membrane, resulting in a low NPo. This result shows that, similar to observations in AT1R transgenic and Dahl spontaneous hypertensive rat models, AC1903 protects podocytes from PAN-induced injury when administered systemically by reducing TRPC5 channel activity.

**Figure 2.**
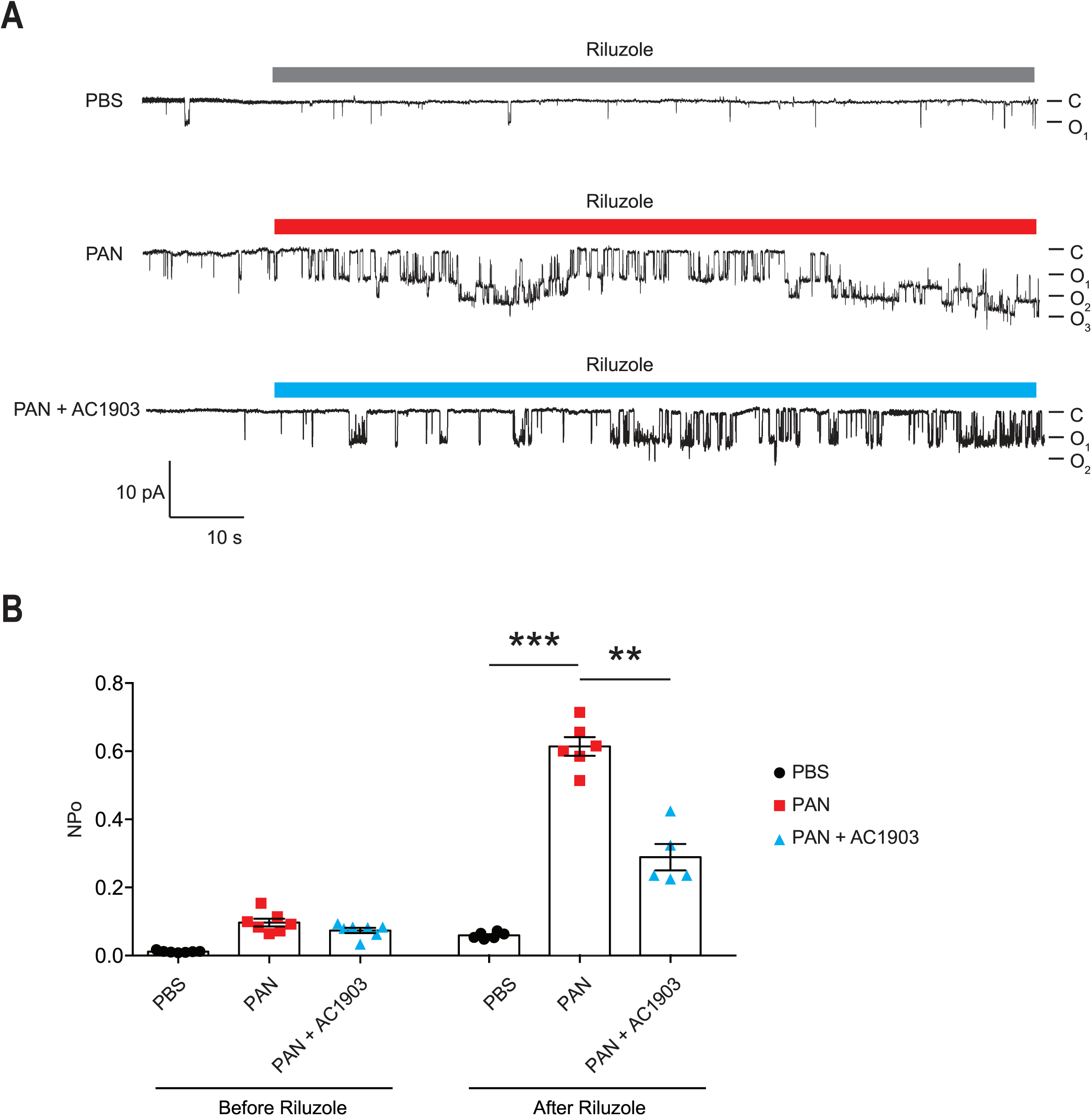
Single-channel recordings from acutely isolated glomeruli show that systemic treatment with AC1903 abrogates TRPC5 activity. (**A**) Representative TRPC5 single-channel current traces from PBS, PAN and PAN + AC1903 treated rats in response to TRPC5 channel agonist Riluzole (30 µM). (**B**) Quantification of TRPC5 single-channel activity by analysis of NPo values. *p < 0.05, ***p < 0.001.

### Human iPSC-derived podocytes express functional TRPC5 channels

Generating human podocytes in 2D cultures offers a unique opportunity to conduct mechanistic studies *in vitro*. To generate human iPodos, we adapted a previously published three-step protocol to induce differentiation into intermediate mesoderm, then into nephron progenitors, and finally, into mature podocytes^30^. Mature iPodos exhibited typical *in vitro* mature podocyte morphology characterized by a large and flat cell body with a dense nucleus that resembled mouse and human immortalized podocytes^31, 32^. The iPodos from this protocol expressed the major podocyte markers including podocin, synaptopodin, alpha actinin-4, and WT-1^30^.

We performed patch clamp electrophysiology, the gold standard in measuring ion channel activity, using iPodos, 12 to 14 days after induction. For whole-cell patch clamp recordings, a single iPodo was identified and the glass pipette was moved to the center of the cell body to provide a more effective Giga-seal (Fig. 3A). Upon successful achievement of the whole-cell configuration, a strong outwardly rectifying current was observed upon application of a voltage ramp protocol, which decreased gradually within 30 seconds of perfusion (Fig. 3B). Englerin A, a compound known to be a nanomolar activator of TRPC4 and TRPC5, was applied once the baseline became stable. Large outward and inward currents were induced by 100 nM Englerin A, which could be blocked by AC1903 (Fig. 3B-C). The inhibitory effect of AC1903 was more prominent at negative potentials confirming that the baseline outwardly rectifying current did not correspond to a TRPC5 conductance (Fig. 3D). These data provide the first evidence that human podocytes express functional TRPC5 channels at baseline, without additional manipulation, indicating that TRPC5 channels may play a role in human podocyte physiology.

**Figure 3.**
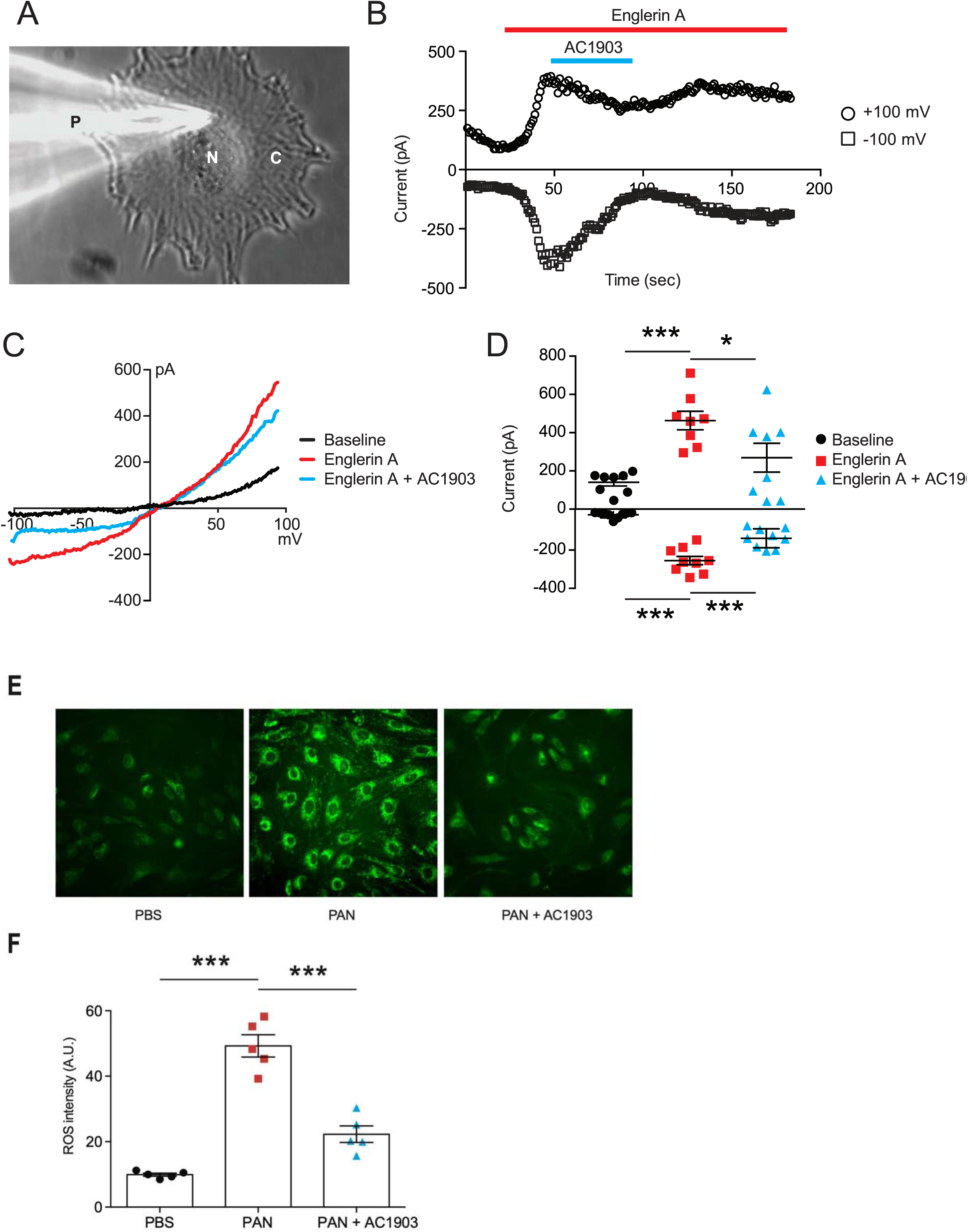
Functional TRPC5 channel activity blocked by AC1903 reduces cytosolic ROS and protects cytoskeletal proteins in PAN-treated iPodos. (**A**) Representative image of a human iPodo patch clamp recording in the whole-cell configuration. P: Glass pipette, N: iPodo nuclear, C: iPodo cytosol. (**B**) Representative diary plots of whole-cell currents from iPodos in response to TRPC5 channel agonist Englerin A (100 nM) in the absence or presence of TRPC5 channel inhibitor AC1903 (30 µM). Currents shown are from +100 mV and −100 mV of a ramp protocol. (**C**) Representative TRPC5 channel current-voltage (I-V) curves from iPodo whole-cell recording. (**D**) Statistical analysis of I-V curves from iPodos treated with Englerin A in the absence or presence of AC1903. (**E**) Representative cytosolic ROS images in iPodos after 24-hour treatment with PAN (150 µg/mL) with or without AC1903 (30 µM). (**F**) Statistical analysis of the ROS signal intensities.

Previous studies have indicated that TRPC5 activity is a major cause for podocyte injury in various rodent *in vitro* and *in vivo* models^16, 33^. To determine whether inhibition of TRPC5 channel activity is protective in human podocytes, we investigated the effect of the TRPC5 channel inhibitor AC1903 on PAN-treated mature human iPodos. We hypothesized that PAN treatment would cause human iPodo injury by triggering the Rac1-TRPC5 feedforward pathway and increasing ROS^16^. In support of this hypothesis, incubation with PAN for 24 hours significantly increased iPodo intracellular ROS levels, which were reduced by co-treatment with AC1903 (Fig. 3E-F). These results suggest that inhibition of TRPC5 channels by the small molecule AC1903 can protect human iPodos from PAN-induced ROS generation. Previous experiments in mouse podocytes have shown that AC1903 blocks ROS generation induced by angiotensin II (AngII) suggesting that both mouse immortalized podocytes^16^, and now human iPodos, support a role for TRPC5 in podocyte biology and disease pathophysiology. In summary, using iPodos, we demonstrated the presence of active TRPC5 channels, blocked them with AC1903, and measured downstream reduction of ROS, the sequela of PAN-mediated Rac1-TRPC5 activation.

### TRPC5 inhibition preserves podocin, synaptopodin, and nephrin abundance in PAN-treated human kidney organoids

To evaluate the effect of AC1903 in human kidney tissue, we took advantage of the human iPSC-derived kidney organoid model. Organoids contain self-organized nephrons composed of early glomerular structures connected to tubular cells including proximal tubules, loops of Henle and distal tubules. These 3D organoids thus hold the potential to be excellent *in vitro* models for preclinical drug testing, because they allow simultaneous monitoring of drug effects on multiple kidney cell types^34-36^. Using immunofluorescence imaging, we found that PAN treatment reduced nephrin, podocin and synaptopodin, but not WT1 expression levels (Fig. 4 and Supplemental Fig. 2). Co-treatment with AC1903 preserved podocyte cytoskeletal proteins, as observed in iPodos and PAN rats. Taken together, our data suggest that inhibition of TRPC5 channel activity can protect human podocytes from PAN-induced injury in an *in vitro* 3D model of the human kidney.

**Figure 4.**
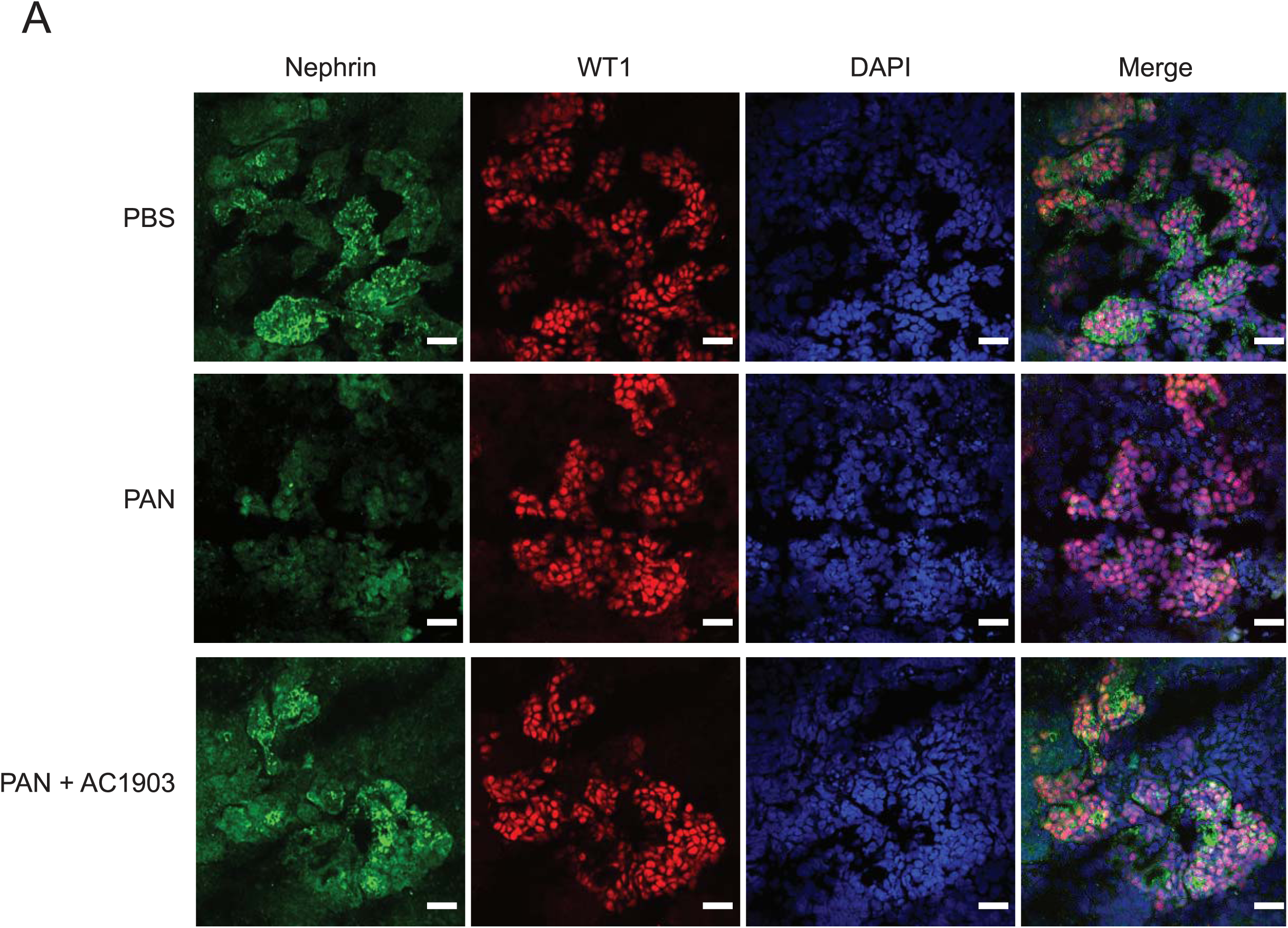
A blocker of the Rac1-TRPC5 pathway protects podocytes from injury in human kidney organoids. (**A**) Representative immunostaining images of podocyte cytoskeletal and marker proteins, nephrin (Green) and WT1 (Red) in PBS, PAN and PAN + AC1903 treated human kidney organoids. Scale bar 20 µm.

## Discussion

The majority of studies investigating podocyte biology *in vitro* have relied on immortalized cell lines. Although these cell lines express podocyte markers such as podocin, synaptopodin, nephrin, and WT-1, and respond to many stimuli, they are prone to de-differentiation, resulting in variability. Moreover, immortalized cell lines fall short of recapitulating the crosstalk and interactions between podocytes and other cell types of the kidney. Following successful generation of 2D human podocyte cultures (iPodos) and 3D kidney organoids *in vitro* from human iPSCs, we showed that PAN treatment causes elevated ROS in iPodos, and podocyte injury in kidney organoids, through the activation of the Rac1-TRPC5 pathway and disruption of cytoskeletal protein dynamics. We demonstrated the podocyte-protective effects of small-molecule inhibitors of TRPC5 channels in human podocytes and kidney organoids, establishing these systems as reproducible, human-specific tools to study podocyte-associated kidney disease^30, 34-36^.

Our previous data demonstrated that a specific TRPC5 small-molecule inhibitor, AC1903, can rescue podocytes and attenuate the progression of kidney diseases in angiotensin II type 1 receptor transgenic, and spontaneous hypertensive rat models^16, 33^. These findings provided a mechanistic rationale for therapeutically targeting TRPC5 channels in the treatment of progressive chronic kidney diseases. In this study, we generated human iPSC-derived podocytes and used patch clamp electrophysiology to demonstrate their response to the TRPC4/5 channel agonist, Englerin A, which was blocked by the TRPC5-selective inhibitor AC1903. To our knowledge, this is the first demonstration that human podocytes express functional TRPC5 channels, further strengthening the notion that these channels play an important role in progressive kidney diseases.

Prior work has shown that Rac1 activation is a nodal event in a spectrum of glomerular diseases, while inhibition of Rac1 activity ameliorates podocyte injury in response to various noxious stimuli^14, 37-40^. In this study, we investigated the contribution of the Rac1-TRPC5 feedforward pathway in a PAN-induced nephrosis rat model. A single dose of PAN was sufficient to induce podocyte injury and proteinuria in rats within a week, which was consistent with the results from PAN-treated human podocytes and kidney organoids. Inhibition of TRPC5 by AC1903 was sufficient to protect podocyte cytoskeletal proteins and suppress proteinuria in PAN-induced nephrosis rats within a week. The accelerated timeline of injury in PAN-induced nephrosis increases the translatability of this model in the context of pre-clinical studies, especially in comparison to AT1R transgenic and Dah1 Salt-sensitive spontaneous hypertensive rat models which require at least a month for podocyte injury and proteinuria to be established.

Taken together, these data indicate that inhibition of TRPC5 channel activity blocks the Rac1-TRPC5 feedforward pathway and protects podocytes from PAN-induced injury. Our data also highlight the utility of human iPodos, kidney organoids, and the PAN-induced nephrosis rat model as useful tools for the preclinical development of TRPC5 channel inhibitors. This diverse set of validated models spans both human *in vitro* systems conducive for mechanistic studies and experimentally tractable *in vivo* disease models with physiological readouts. In sum, this study bolsters the human relevance and scientific rationale for a TRPC5-targeted podocyte-protective strategy.

## Acknowledgements

This work was supported by NIH/NIDDK grants DK103658, DK099465 and DK095045 (A.G.).

## Methods

### Animals

Wild-type Sprague-Dawley rats (Male, 4-5 weeks, Charles River) were housed under a controlled environment with a 12-hour light-dark cycle and access to food and water ad libitum. All animal experiments were performed in accordance with the guidelines established and approved by the Animal Care and Use Committee at Brigham and Women’s Hospital, Harvard Medical School. After wild-type Sprague-Dawley rats were acclimated for a week in the BWH CCM animal facility. A single dose of puromycin aminonucleoside (50 mg/kg, PAN group) was given i.p. to rats to induce nephrosis, and PBS was given as control. Following the PAN injection, vehicle or AC1903 (50 mg/kg) was administered twice daily (at 9 am and 9 pm) for seven days. 24-hour urine albumin levels were measured on day 0, 3 and 7. Rats were euthanized after the metabolic collection on Day 8. Both kidneys were collected for downstream experiments. In most cases, one kidney was used for acute glomeruli isolation and the other was stored in −80°C for subsequent experiments. In combination, we have studied 34 rats (PBS group n = 6, PAN group n = 15, PAN + AC1903 group n = 13).

### Chemical preparation and IP administration

All chemicals were purchased from Sigma-Aldrich unless described otherwise. AC1903 was synthesized, purified and prepared by C.H. as previously published^16^. Immediately prior to injections, AC1903 solution was placed on a heated shaker at 48°C and 800 rpm. Vehicle was prepared in the same fashion. Injection amount was determined by body weight (2mL vehicle/compound per kg body weight). Body weight was measured at the time of injection.

### Metabolic collection and urine albumin assay

Rats were housed individually in a metabolic cage supplied with adequate amounts of food and water. Urine was collected into a 50 mL Falcon tube for 24 hours. Total urine volume was measured and then centrifuged at 3,200x g for 10 min at 4°C. Albumin quantification was done according to our previously published protocol^16^. Coomassie Brilliant Blue stained gels of urine samples were quantified by densitometry with albumin standards using Image J software.

### Human iPSC culture

Human Episomal iPSC Line (ThF) (ThermoFisher, #A18945) was maintained in mTeSR1 medium (Stem Cell Technologies, #85870) in T25 flasks pre-coated with Matrigel (Stem Cell Technologies, #354277). Cells were passaged using Gentle Cell Dissociation Reagent (Stem Cell Technologies, #7174). iPSCs were confirmed to be karyotype normal and maintained below passage 10 and all the cell lines were routinely checked and were negative for mycoplasma.

### Differentiation into human iPSC-derived podocytes (iPodos)

Human iPSC-derived podocytes (iPodos) were generated using the cited protocol with a few modifications ^30^. A total number of 3.75 × 10^5^ ThF human iPSCs were seeded in a Matrigel-coated T25 flask in mTeSR1 medium (Stem Cell Technologies, #85870) with ROCK inhibitor, Y-27632 (10 µM, Stem cell Technologies, #72304). After 24h cells were treated with a 1:1 mixture of DMEM/F12 + GlutaMAX (Life Technologies, #10565-018) and Neurobasal media, supplemented with N2 and B27 (Life Technologies, #21103049), CP21R7 (1 µM, Cayman Chemical, #20573), and BMP4 (25 ng/mL, Peprotech, #AF-120-05ET), for three days. On day four, the medium was replaced with STEMdiff APEL2 medium (Stem Cell Technologies, #05270) supplemented with FGF9 (200 ng/mL, Peprotech, #100-23), BMP7 (50 ng/mL, Peprotech, #120-03), and Retinoic Acid (100 nM, Sigma-Aldrich, #R2625) for two days. On day six, cells were dissociated with Accutase (Stem Cell Technologies, #07920) and 2 × 10^5^ cells were seeded on Type I Collagen-coated 6-well dishes and cultured until day fourteen in DMEM/F12+GlutaMAX medium supplemented with 10% FBS (Life Technologies, #16140071). Vitamin D3 (100 nM, Tocris Bioscience, #4156), and Retinoic Acid (100 µM, Sigma-Aldrich, #R2625) were added every other day. Cells were fully differentiated and ready to use from Day 12 to Day 14.

### Kidney organoid differentiation

Kidney organoids were generated using a previously described protocol ^34^ with slight modifications. A total number of 3.75 × 10^5^ ThF iPSCs were plated in a T25 flask in the mTeSR1 medium with ROCK Inhibitor Y-27632 (10 µM, Stem cell Technologies, #72304). After 24 hours, cells were treated with CHIR99021 (8 µM, R&D systems, #4423/10) in the STEMdiff APEL2 medium (Stem Cell Technologies, #05270) for four days, followed by recombinant human FGF-9 (200 ng/mL, Peprotech, #100-23) and heparin (1 µg/mL, Sigma-Aldrich, #H4784) for an additional three days. At day seven, cells were dissociated into single cells using AccutaseTM (Stem Cell Technologies, #07920). 5 × 10^5^ cells were pelleted at 350x g for 2 min and transferred onto a 6-well transwell membrane (Stem Cell Technologies, #3450). Pellets were incubated with CHIR99021 (5 µM) in the APEL2 medium for one hour at 37°C. Then the medium was changed to the APEL2 medium with FGF-9 (200 ng/mL) and heparin (1 µg/mL) for an additional five days, and an additional two days with heparin (1 µg/mL). Medium was changed every other day. The organoids were maintained in APEL2 medium with no additional factors until day 25. Then kidney organoids were treated with PBS, PAN (150 µg/mL) with or without AC1903 (30 µM) for 72 h before the downstream experiments.

### ROS assay

Human iPSC-derived podocytes (iPodos) were treated with either PBS, PAN (150 µg/mL), or PAN with 30 µM AC1903 for 24 h. Intracellular production of ROS was measured using a cell-permeable fluorescent ROS indicator (Invitrogen, #C10444) following the official protocol. Briefly, cells were incubated with 5 µM CellRox Green at 37°C for 30 min in Hanks’ balanced salt solution (ThermoFisher Scientific, #14025092). Cells were then washed with PBS and fixed with 4% PFA. Fluorescence images were taken under a confocal microscope Olympus FV-1000. ROS signal intensities were quantified using ImageJ software.

### Rat kidney immunofluorescence

Rats were euthanized and perfused with PBS. The kidney was quickly removed and cut into half. One half was flash frozen in liquid nitrogen, and the other was fixed in 4% PFA overnight and stored in PBS for follow-up experiments. For immunofluorescence, Kidney tissues were sectioned at 6 µm thickness and blocked with 3% BSA at room temperature for 1 h. The rabbit anti podocin, guinea pig anti synpo, goat anti Nephrin, and rabbit anti WT-1 antibodies were used at a dilution of 1:200. The Alexa goat anti rabbit and guinea pig IgG 488 and Alexa donkey anti goat IgG 594 were used at a dilution of 1:200. Fluorescence images were taken with a confocal microscope Olympus FV-1000.

### Electrophysiology

iPodo whole-cell patch clamp was performed using an Axopatch 700B and Digidata 1550A (Molecular Devices). Bath solution contained (in mM) 135 CH_3_SO_3_Na, 5 CsCl, 2 CaCl_2_, 1 MgCl_2_, 10 HEPES, and 10 glucose adjusted with NaOH to pH 7.4. Pipette solution contained (in mM) 135 CH_3_SO_3_Cs, 10 CsCl, 3 MgATP, 0.2 EGTA, 0.13 CaCl_2_, and 10 HEPES adjusted with CsOH to pH 7.4. For glomerular single-channel recording, acutely isolated glomeruli were prepared as previously published ^16^. Single-channel recordings were carried out using an Axopatch 200B and Digidata 1550A (Molecular Devices). Bath and pipette solutions for glomerular single-channel recording contained (in mM) 135 CH_3_SO_3_Na, 5 CsCl, 2 CaCl_2_, 1 MgCl_2_, 10 HEPES, and 10 glucose adjusted with NaOH to pH 7.4. Once the inside-out configuration was achieved, the bath solution was replaced by an intracellular solution, containing (in mM) 135 CH_3_SO_3_Cs, 10 CsCl, 3 MgATP, 0.2 EGTA, 0.13 CaCl_2_, and 10 HEPES adjusted with CsOH to pH 7.4. Patch pipettes, with a resistance of 4-6 MΩ, were prepared using a two step-protocol (Sutter Instrument, P-97). Pipettes were fire-polished before use with a microforge (Narishige, MF-9). For glomerular single-channel recording, data was acquired at 10 kHz sampling frequency, and filtered with low-pass filtering at 1 kHz. Holding membrane potential was at −60 mV. Single-channel analysis was carried out using Clampfit 10.4 software (Molecular Devices). NPo were analyzed for 10 sec before and after the application of TRPC5 agonist riluzole (Sigma, R116).

### Statistical analysis

All the data were presented as Mean ± SEMs unless described otherwise. Microsoft Office Excel, Origin 6.0 and Graphpad Prism 6 were used for statistical analysis and creation of the graphs. For statistical analysis of differences, an unpaired t-test and a one-way ANOVA followed by Bonferroni or Tukey Correction were used. P value less than 0.05 was considered to be significant.

---

**Supplementary Figure 1.**
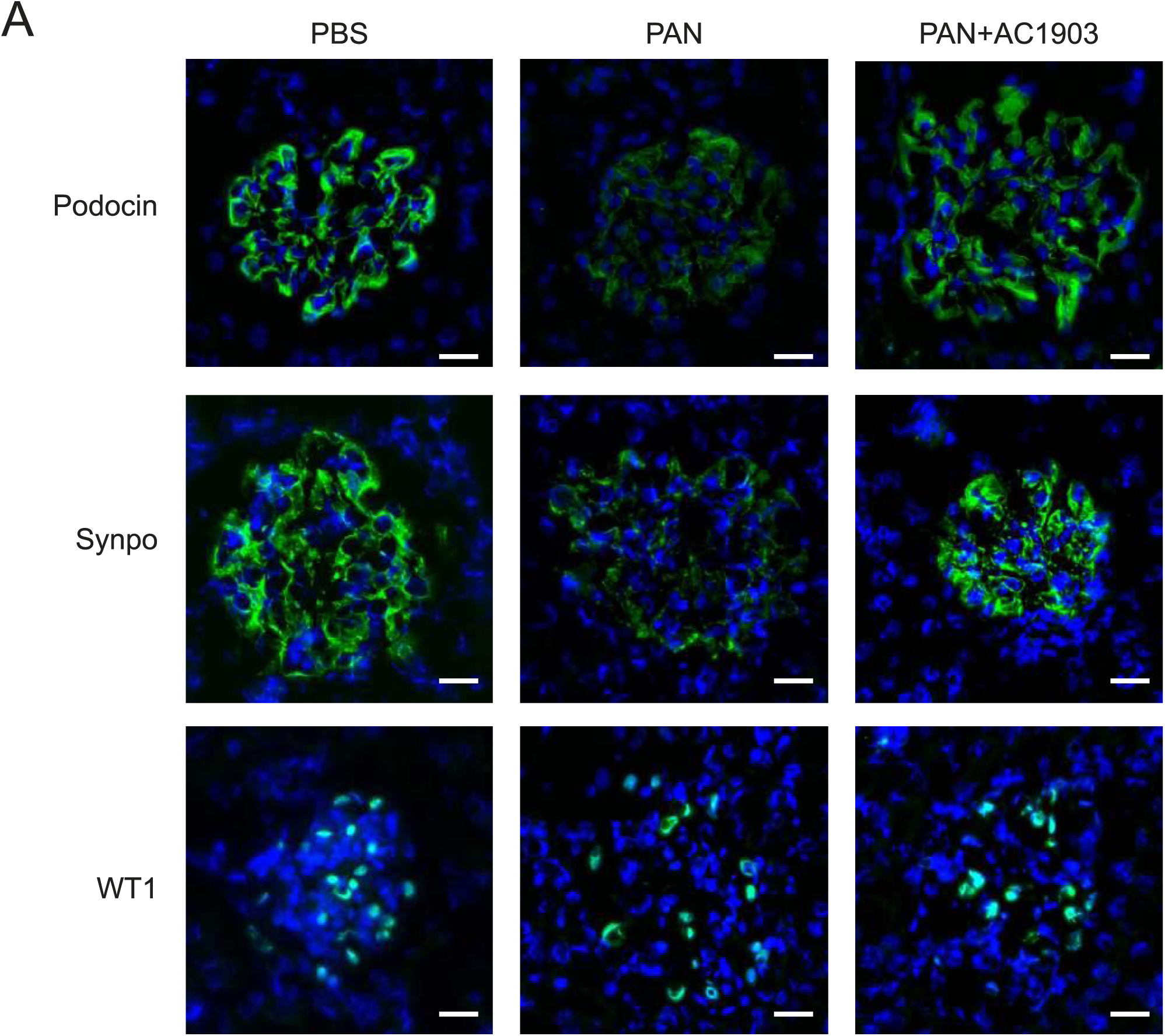
Inhibition of TRPC5 channel activity protects podocytes in PAN rats. (**A**) Representative immunostaining images of podocyte cytoskeletal and marker proteins podocin, synaptopodin, and WT1 from PBS, PAN and PAN +AC1903 treated rats on day 7. Scale bar 20 µm.

**Supplementary Figure 2.**
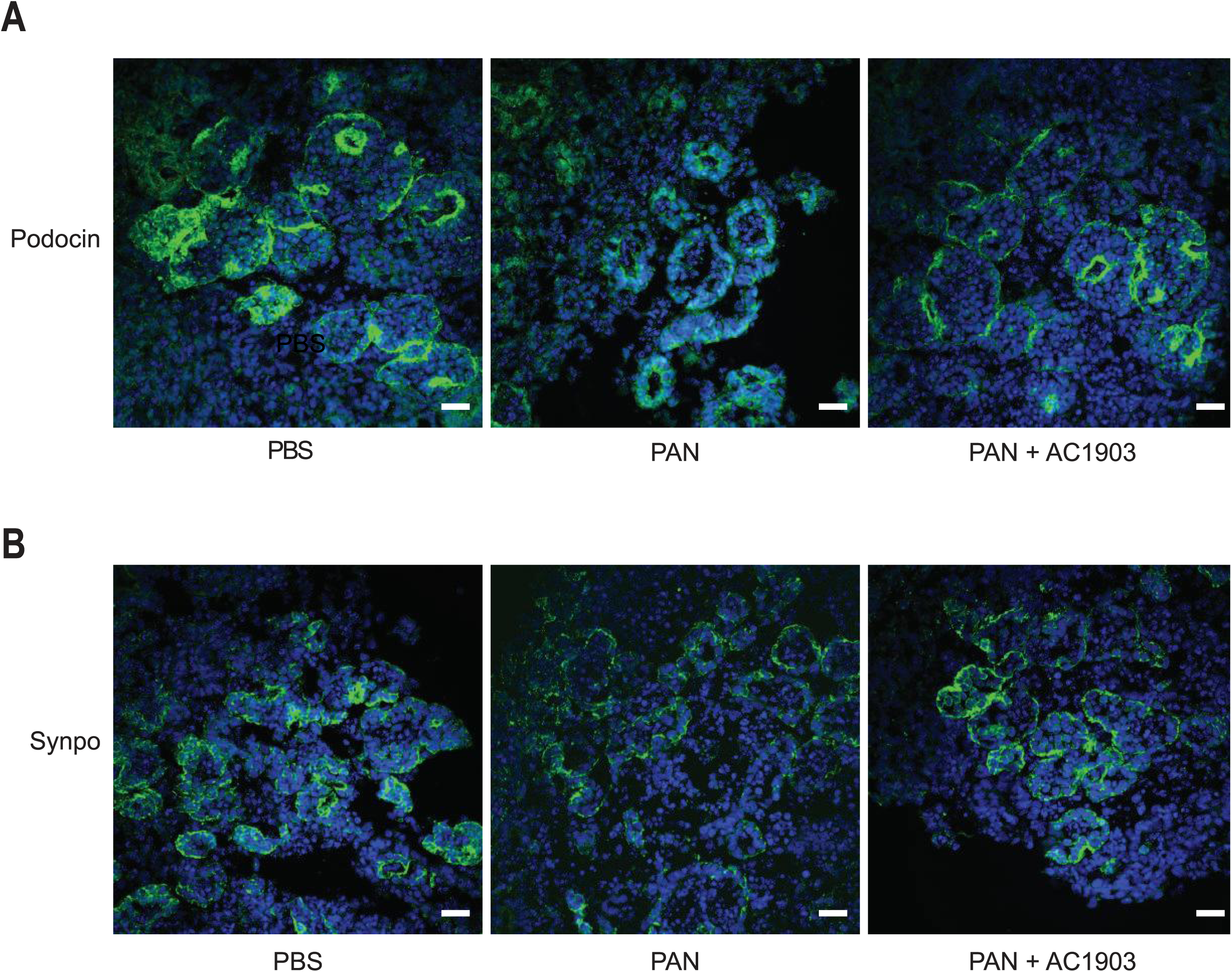
Inhibition of TRPC5 activity protects podocin and synaptopodin in human kidney organoids treated with PAN. (A, B) Immunostaining for podocin (A) and synaptopodin (B) in PBS, PAN and PAN + AC1903 treated kidney organoids. Scale bar 20 µm.

